# Chimeras Link to Tandem Repeats and Transposable Elements in Tetraploid Hybrid Fish

**DOI:** 10.1101/088070

**Authors:** Lihai Ye, Xiaojun Tang, Yiyi Chen, Li Ren, Fangzhou Hu, Shi Wang, Ming Wen, Chun Zhang, Ming Tao, Rurong Zhao, Zhanzhou Yao, Shaojun Liu

## Abstract

The formation of the allotetraploid hybrid lineage (4nAT) encompasses both distant hybridization and polyploidization processes. The allotetraploid offspring have two sets of sub-genomes inherited from both parental species and therefore it is important to explore its genetic structure. Herein, we construct a bacterial artificial chromosome library of allotetraploids, and then sequence and analyze the full-length sequences of 19 bacterial artificial chromosomes. Sixty-eight DNA chimeras are identified, which are divided into four models according to the distribution of the genomic DNA derived from the parents. Among the 68 genetic chimeras, 44 (64.71%) are linked to tandem repeats (TRs) and 23 (33.82%) are linked to transposable elements (TEs). The chimeras linked to TRs are related to slipped-strand mispairing and double-strand break repair while the chimeras linked to TEs are benefit from the intervention of recombinases. In addition, TRs and TEs are linked not only with the recombinations, but also with the insertions/deletions of DNA segments. We conclude that DNA chimeras accompanied by TRs and TEs coordinate a balance between the sub-genomes derived from the parents which reduces the genomic shock effects and favors the evolutionary and adaptive capacity of the allotetraploidization. It is the first report on the relationship between formation of the DNA chimeras and TRs and TEs in the polyploid animals.

## Introduction

It is certain that chimeric genes are the results of recombinations between DNA sequences from various sources with various functions. Recombination can provide the raw materials for biological evolution and it enables the reconstruction and rearrangement of genome to eliminate deleterious mutations. Homology is the main factor driving the recombination of any two sequences (Gaeta and Chris 2010). Researches show that recombination can occur among repeats within the same chromosome, on homologous chromosomes (Jelesko *et al.* 2004) and even among nonhomologous chromosomes that share some degree of homology (Mezard *et al.* 2007). Recombination is closely linked with DSB (double-strand break) repair and it is useful for the restoration of broken replication forks (Heyer and Ehmsen 2010). What we concern is how the recombinations occur and what causes the chimeras in polyploids. Recombination in allopolyploids plants may occur ectopically among paralogous (Jelesko *et al.* 2004) or homoeologous (Qi *et al.* 2007) sequences because of a lack of diploid pairing fidelity. In newly formed allopolyploids of *Brasssica napus*, homoeologous recombinations are deeply entwined with reciprocal exchanges and gene conversions, and are responsible for many of genetic changes (Gaeta and Chris 2010). In addition, nonreciprocal homoeologous exchanges have occurred throughout polyploid divergence and speciation in allopolyploid cotton *(Gossypium)* (Salmon *et al.* 2010). Considering polyploidy increases the number of duplicated sequences resident in the genome, homology-dependent chimeras could also be increased correspondingly. However, there is no doubt that there are some other factors that could cause chimeras in polyploid.

Although our laboratory has conducted research on chimeras in allotetraploid fish (Liu *et al.* 2016), the repetitive elements adjacent to those chimeras were not even noticed. There are investigations implicate that repetitive elements play a role in the generation of chimeras (Jiang *et al.* 2004; Kapitonov *et al.* 2006). Repetitive elements have historically been called “junk DNA”; however, they are important forces for recombination and are essential for genome evolution (Shapiro and Sternberg 2005). Major repetitive elements include tandem repeats (TRs) and transposable elements (TEs). TRs comprise repeat units that are directly adjacent to each other, and because of their particular structure, the DNA sequences around them are unstable (Bichara *et al.* 2006). Two mechanisms have been proposed to explain this instability: unequal recombination and DNA polymerase slippage (Debrauwere *et al.* 1997). There are differences in content of TRs among different species (Toth and Gaspari 2002), among different chromosomes within the same species (Katti *et al.* 2001), and between coding and non-coding regions (Catasti *et al.* 1999; Cox and Mirkin 1997). The variable TRs can change the gene structure, or even influence gene expression and evolution (Gemayel *et al.* 2010). The transposable element is a large member of repetitive DNA, which can transfer to new genome sites and reproduce themselves during the process. They intersperse in the genomes of plants and animals, occupying a large proportion; thus, they may not only alter individual gene structure, but also the genome structure and function (Bennetzen 2000). In addition, they have vital influences on the structural diploidization of genomic DNA (Lim *et al.* 2007; Bruggmann *et al.* 2006) and the early stage of the evolution of allopolyploids (Oliver and greene 2009). Undoubtedly, they are an important component of an organism’s genome and are irreplaceable for biological evolution, genetic heredity and variation. Thus, repetitive elements provide a great platform for the study of genetic variation in polyploids.

Polyploids are organisms with three or more complete sets of chromosomes, and many plants and animals have experienced polyploidization one or more times during their evolution (Song *et al.* 2012; Masterson 1994). The essence of allopolyploid formation is the fusion of genomes of two species, which develops into a hybrid whose phenotype and genotype are different from the parental species. However, the key mechanism of their genome reconstruction is homoeologous pairing and recombination (Gaeta and Chris 2010). Homoeologous recombination is observed in allopolyploids of *Lolium multiflorum-Festuca* by genomic in situ hybridization (Zwierzykowski *et al.* 1998). In the genome of resynthesized *Brasssica napus* allopolyploids, many genomic changes are detected, such as recombinations, deletions, replication, and translations (Gaeta and Chris 2010). But the allopolyploid formation is very rare in animals and even less reports has focused on recombination in animal polyploids. The allotetraploid fish hybrid (4nAT) is created by our laboratory via an artificial intergeneric cross between red crucian carp (RCC) *(Carassius auratus* red var., ♀, 2n = 100) and common carp (CC) *(Cyprinus carpio* L., ♂, 2n = 100) (Liu *et al.* 2001) and to date, the 25 ^th^ generation of 4nAT has been formed through successive self-breeding. The 4nAT represents the foundation for the speciation of a new tetraploid fish and provide perfect materials for studying the effects of distant hybridization and polyploidization on genome evolution. However, the allotetraploids has two genomes inherited from diploid progenitors, what is its structure? How do the surviving tetraploid hybrids overcome the genomic shock effects caused by the reintegration of two different genomes? These are the questions we seek to answer in the present study. Herein, we construct a bacterial artificial chromosome (BAC) library of allotetraploid offspring and compared the BAC sequences with their parental genomes to analyze the relationship between the mechanisms of genetic changes, such as gene recombination, insertions/deletions (indels) and repetitive elements, which will provide new impetus for research on the genome evolution of polyploids.

## Materials and Methods

### The constructing of 4nAT BAC library

Blood samples (10 ml) from five allotetraploid individuals from the 20^th^ generation are used to construct a BAC library according to a previously described method (Wang *et al.* 2015). The allotetraploid fish are obtained from the Engineering Research Center of Polyploid Fish Breeding at Hunan Normal University (Changsha, China) and the ploidy of five samples is confirmed by metaphase chromosome assay of kidney cells. The genomic DNA of allotetraploids is digested with the restriction enzyme *Hind* III, ligates into the CopyControl pCC1BAC vector, and then transforms into *Escherichia coli DH10B*. The recombinant bacterial clones are screened through blue-white screening. Nineteen BAC clones are picked randomly and sequenced by Shanghai Majorbio Bio-pharm Technology Co. Ltd. The sequencing and assembly of the 19 BAC clones are performed by Illumina next-generation sequencing technology and PacBio RS platform.

### Data processing

NCBI-VecScreen is used to remove the vector sequences from the obtained BAC sequences (http://www.ncbi.nlm.nih.gov/tools/vecscreen/) and RepeatMasker (http://www.repeatmasker.org/) is used to calculate the types and contents of repetitive elements in the allotetraploids BAC sequences, using *Danio rerio* as the reference organism. We compare the BAC sequences that include repetitive elements with the genomes of both progenitors (using NCBI-BLAST-2.3.0+) and homologous sequences are identified with an *E*-value <1e-5 and similarity >95%. The maternal progenitor genome sequencing has been completed by our laboratory together with Yunnan University and the database has yet to be published (http://rd.biocloud.org.cn/). The genome of paternal progenitor can be searched from the NCBI database (http://www.ncbi.nlm.nih.gov/) and the Common Carp Genome Database (http://www.carpbase.org/).

### Models of chimeras based on BAC analysis

We calculate the number of chimeras DNA link to repetitive elements in the allotetraploid BAC sequences and then survey the types and presence of the repetitive elements in the parental homologous sequences. Almost all the recombinations (except for the *FN1* gene loci) detected in this study are liked with repetitive elements and they can be divided into four composite patterns according to the presence of repetitive elements in the parental homologous sequences. The four composite patterns are showed in Figure 1: (a) repetitive elements existing only in homologous sequences of maternal progenitor; (b) repetitive elements existing only in homologous sequences of paternal progenitor; (c) repetitive elements existing in both parental homologous sequences; and (d) repetitive elements that don’t exist in either parental homologous sequence.

**Figure 1.**
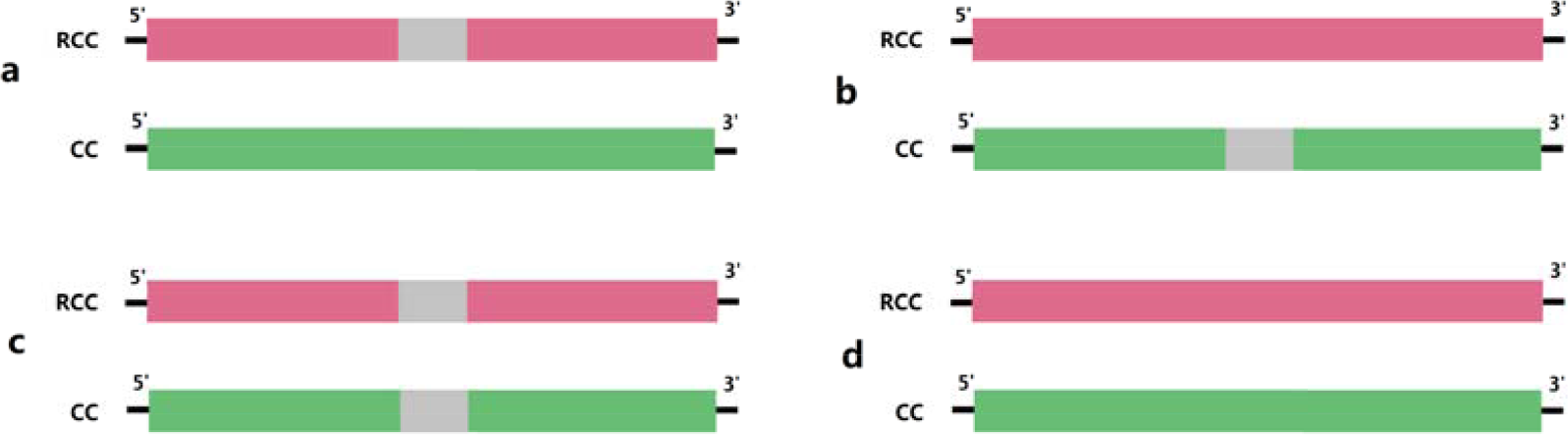
Four composite patterns of parental homologous sequences. The above four composite patterns are divided according to the existent of repetitive element in homologous sequences of parental progenitors.

### Significance difference on statistical analysis

To better explore the mechanism of recombination that resulted from repetitive elements, significance tests of the recombination rates of situations a, b, c and d are conducted. In addition, we analyze the differences among the rates of recombinations related to mono, di, tri, and tetra-nucleotide repeats. The formula involved in the different significance tests is quoted from Du (Du 2003).

## Data availability

The NCBI accession numbers of the 19 full-length BAC clones are: KF758440-KF758444, KJ424354-KJ424362 and KT726912-KT726916. The NCBI accession numbers of the paternal and maternal progenitor homologous sequences are KU508466- KU508484 and KU508485- KU508503 respectively.

## Results

### Statistics of repetitive elements in 4nAT BAC

The BAC library of allotetraploid hybrids has been constructed and 19 full-length BAC clones (NCBI accession numbers: KF758440-KF758444, KJ424354-KJ424362 and KT726912-KT726916) is obtained with a total length of 752,904 bp. The GC content of the 19 BAC clone sequences are range from 34.17% to 41.40%, with an average of 37.31%, and the content of repetitive elements are range from 5.12% to 54.96%, with an average of 16.88%, according to RepeatMasker datasets. The repetitive DNA content of homologous sequences of the maternal red crucian carp and paternal common carp genome are 12.32% and 15.50%, respectively.

We identify 480 TRs, of which 44 (9.17%) are related to recombination (Figure 2A). The results show that the most abundant type of tandem repeats is A/T, whose total number is much larger than that of G/C. The total content of the TRs in 4nAT is negatively correlated with the length of the repeat unit. The statistics confirm that the numbers of chimeras in 4nAT linked to mono, di, tri, and tetra-nucleotide repeats are 11, 24, 2 and 2, respectively. Among the chimeras link to TRs, the numbers of chimeras under composite patterns a, b, c and d are 8, 12, 14 and 10, respectively. The different significance tests of rates of chimeras under the above four situations showed that the rate of recombination under situation c (in which both the homologous sequences of the parental progenitors shared the TR) is much lower than the rates of the other three situations, while the recombination rates of the three other situations showed no remarkable differences (Figure 2B). The different significance tests of the rates of recombinations link to TRs of different repeat unit length show that the recombination rate relate to di-nucleotide repeats is significantly higher than the recombination rates relate to the other three types of TRs and there are no remarkable differences among the three other recombination rates (Figure 2C).

**Figure 2.**
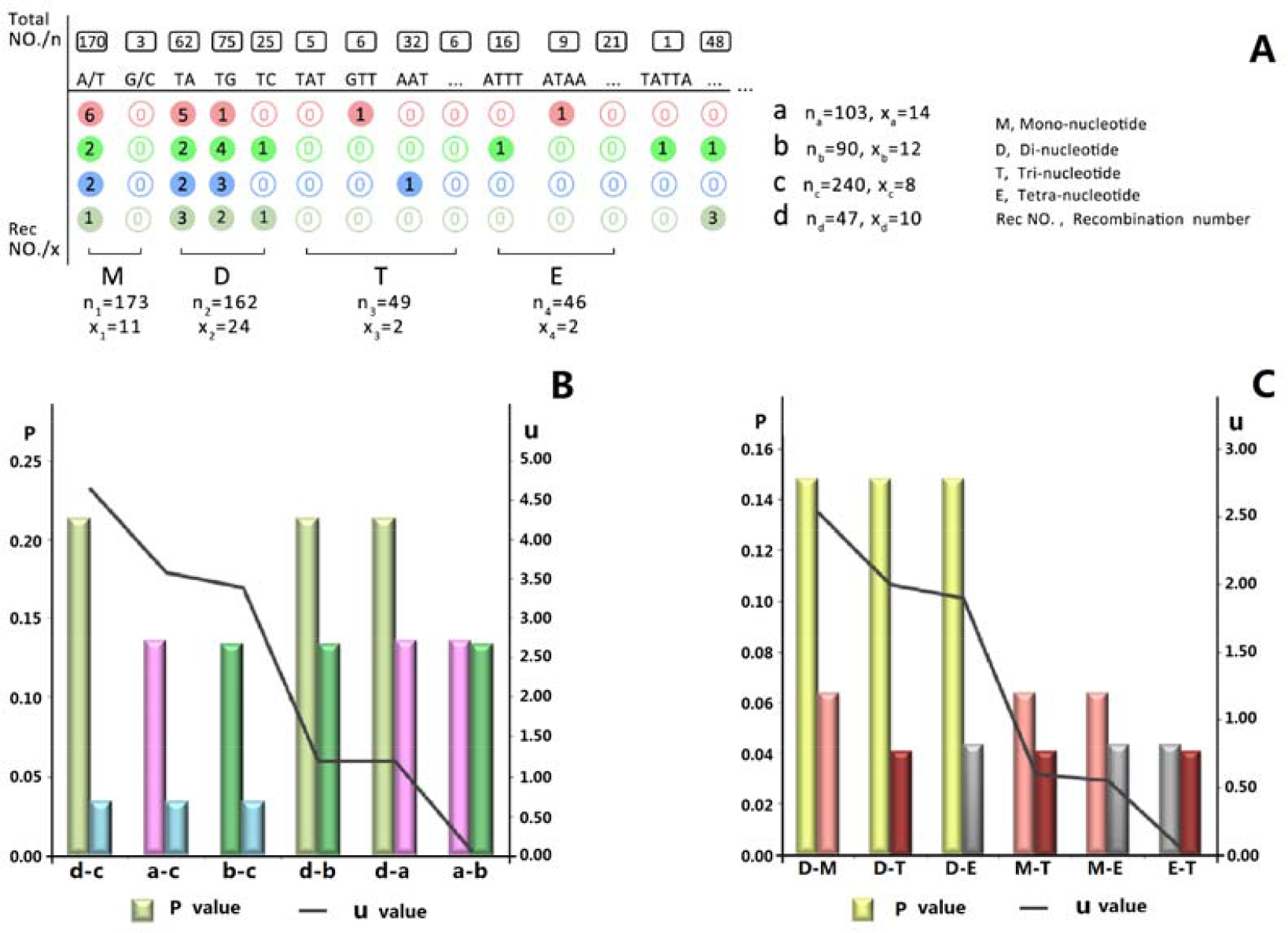
The statistics of tandem repeats. (A) displays the total numbers of tandem repeats and recombination numbers under situations a, b, c and d, respectively. The first and the second apostrophes represent tri-nucleotide TRs and tetra-nucleotide TRs, respectively, and the last apostrophe represents all the penta-nucleotide repeats and TRs with longer units that are not listed here. (B) displays a significance analysis of recombination probabilities among situations a, b, c and d situations, where the u values of a-c, b-c and d-c are greater than u_0.01_ (2.337). (C) displays significance analysis of recombination probabilities among mono-nucleotide, di-nucleotide, tri-nucleotide and tetra-nucleotide repeat loci, where the u values of M-D (mono-nucleotide repeat versus di-nucleotide repeat), D-T (di-nucleotide repeat versus tri-nucleotide repeat), and D-E (di-nucleotide repeat versus tetra-nucleotide repeat) are greater than u_0.05_ (1.645).

We obtain 217 TEs in the 19 BAC sequences, including 139 (64.06%) DNA transposons and 78 (35.94%) retrotransposons. The analysis shows that the distribution densities of TEs in different BAC sequences are different and the maximum density (4nAT-150D6) is 6.9 times greater than the minimum (4nAT-150B4) (Figure 3B). The numbers of chimeras related to TEs under situations a, b, c and d are 2, 2, 18 and 1, respectively (Figure 3A) and the rate of recombination in the above four situations shows no significant differences (u < u0.05 = 1.645, Figure 3C).

**Figure 3.**
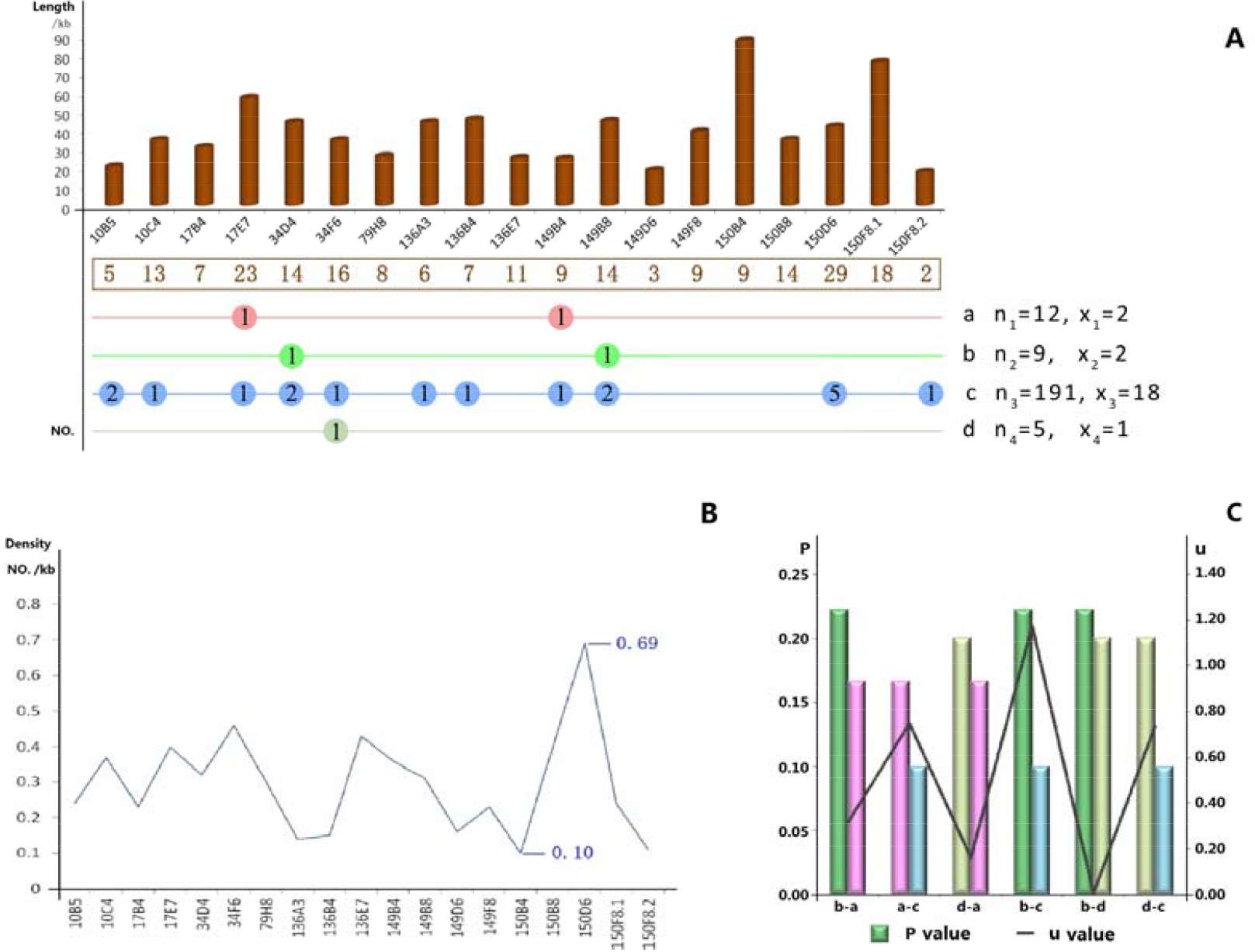
The statistics of transposable elements. (A) displays the total length of every BAC sequence (above the horizontal axis) and the corresponding recombination numbers related to TEs in situations a, b, c and d (below the horizontal axis). (B) displays the density of TEs in every BAC sequences and the biggest and smallest densities were in BACs 4nAT-150D6 (0.69 /kb) and 4nAT-150B4 (0.10 /kb). (C) displays the significance analysis of recombination probabilities among situations a, b, c and d at TE loci: all of the u values are smaller than u_0.05_ (1.645), which means that there were no significant differences among the four situations.

### Models of chimeras

In this study, we confirm that repetitive elements are closely link to recombination. In each of the composite pattern a, b, c and d (Figure 1), there are four chimeras models (Figure 4) that can be identified in allotetraploid offspring according to their genetic source of the two ends: model1, the 5’ end is inherited from RCC while the 3’ end is from CC; model 2, the 5’ end is inherited from CC while the 3’ end is from RCC; model 3, both the 5’ end and the 3’ end are inherited from different scaffolds of RCC; and model 4, both the 5’ end and the 3’ end are inherited from different scaffolds of CC.

**Figure 4.**
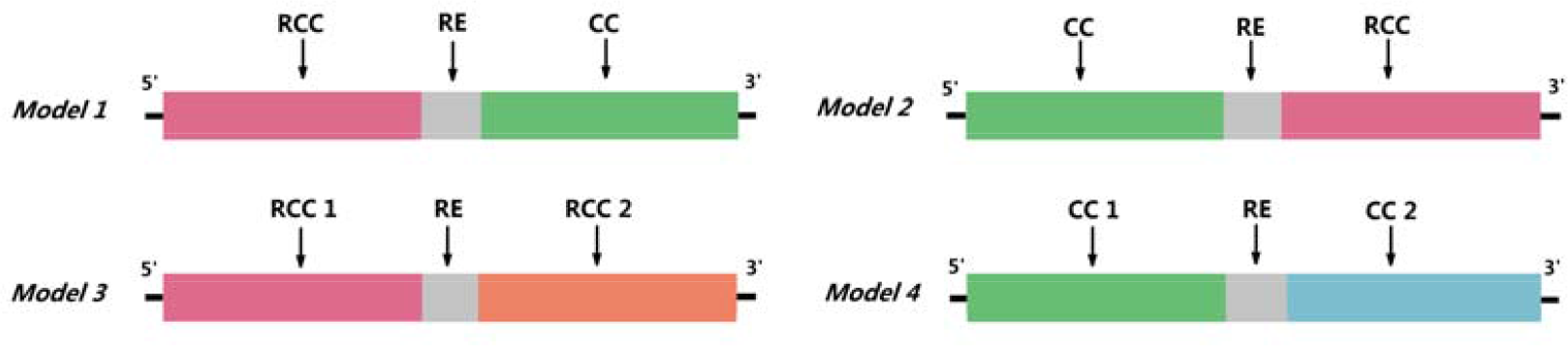
Recombination models linked to repetitive elements. The above four recombination models are divided according to the distribution of the genomic DNA derived from the parents.

### Chimeras link to TRs and TEs

Forty-four (9.17%) out of 480 TR loci are detected to have experienced recombination, and among them, the largest number of recombination sites involve TA repeats, followed by mono-nucleotide A/T repeats. Notably, no DNA chimeras occur at mono-nucleotide G/C repeat loci. Chimeras link to TRs have four models showed in Figure 4, and their specific structures are showed in Figures S1-S4. In addition, there are small pieces of DNA inserted into the recombination locus (see Figures S1, S2 and S4), and according to our statistics, 29.55% of the chimeras have these “little tails” of various lengths.

Twenty-three (10.60%) of 217 TE loci are detected to have experienced recombinations. Among them, 12 are DNA transposons and 11 are retrotransposons. Chimeras related to TEs also have four models showed in Figure 4 and their specific structures are showed in Figures S5-S8. Similarly, there are small DNA segments inserted into the recombination locus too (see Figures S5, S6), and according to our statistics, 24.67% of the chimeras have been inserted with DNA pieces of various lengths.

### Insertion and deletion of TEs

Comparing the BAC sequences with the homologous sequences of both progenitors, insertions/deletions (indels) of TEs are identified in allotetraploids BACs (see Figure 5). By comparing the 4nAT-150D6 BAC sequence to scaffold CC-1513 of paternal genome, we find a retrotransposon Gypsy35-I is inserted into 4nAT-150D6 and a transposon, Mariner-1, is deleted in 4nAT-150D6. Meanwhile, by comparing the BAC sequence to scaffold RCC-200893 of maternal genome, a residue of LINE1 is detected insert into 4nAT-150D6 and a LINE1-2 sequence is deleted in 4nAT-150D6.

**Figure 5.**
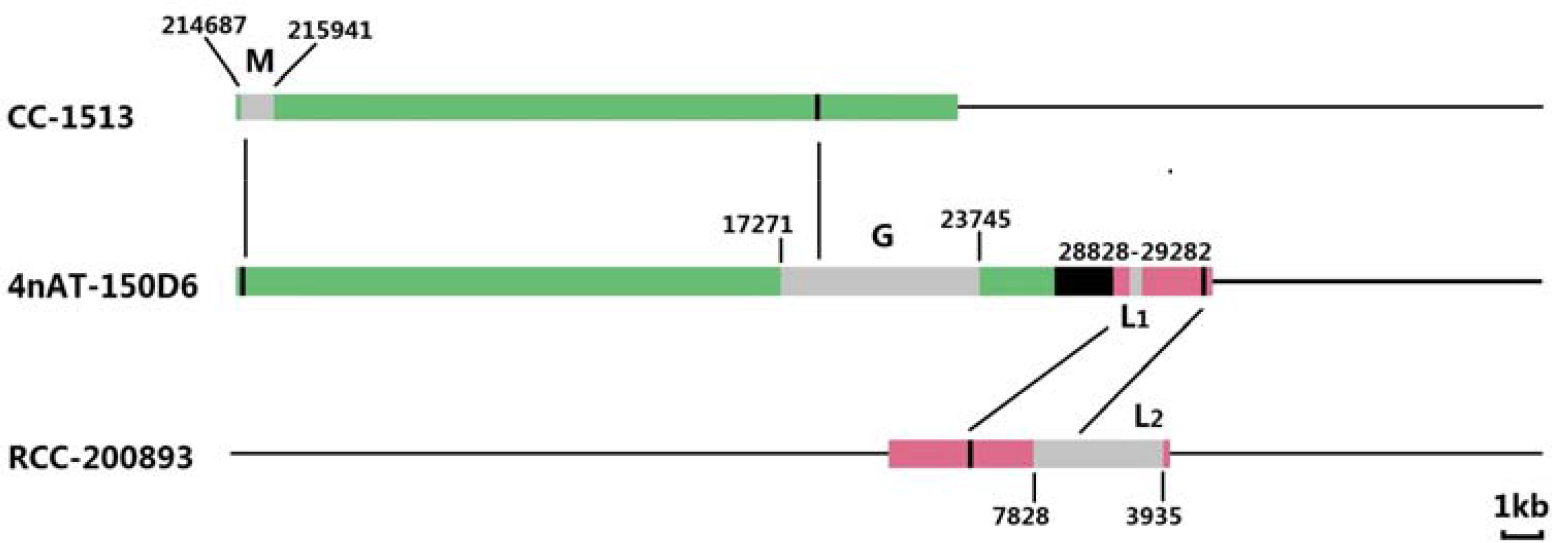
The alignment among BAC 4nAT-150D6 sequence and the parental genomes. CC-1513 and RCC-200893 are the homologous sequences of paternal species and maternal species, respectively. The bars in grey color are transposable elements and M, G, L1 and L2 represent transposon Mariner-1, retrotransposons Gypsy35-I, LINE1 and LINE1-2 respectively. The region in black color in 4nAT-150D6 is shared by both of the parental homologous sequences.

### Chimeras in *FN1* Gene

There is only one chimera detected at non-repetitive element loci among the 19 BAC clone sequences of tetraploid hybrids, which is the *fibronectin 1 (FN1)* gene in BAC 4nAT-10C4. The *FN* gene family encodes fibronectin and is highly conserved in evolution. The *FN1* genes of maternal red crucian carp and paternal common carp have high sequence similarity and some of their coding regions are almost identical; therefore, homoeologous recombination could occur at the *FN1* gene of 4nAT driven by homology. *Fibronectin* is composed of two strands and each strand consists of a chain of repeat unit. So, although *FN1* gene is not repetitive element, it has the similar structure. Figure 6 shows the structure of *FN1* Gene in BAC 4nAT-10C4.

**Figure 6.**
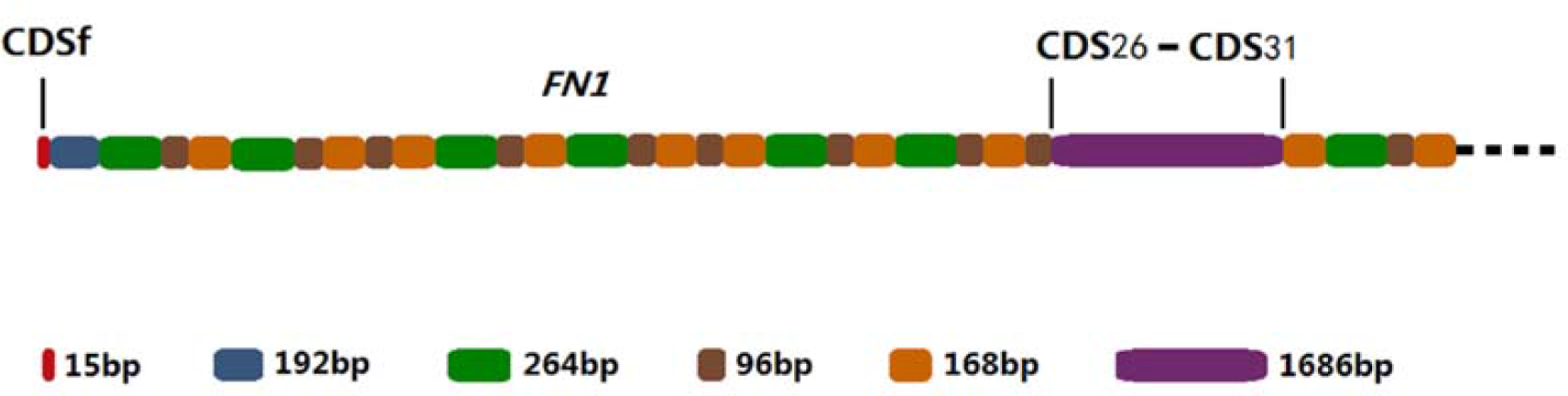
The structure of *FN1* gene in BAC clone 4nAT-10C4. Because of the limited length of BAC clone 4nAT-10C4, *FN1* gene showed here is not complete, contain 35 exons.

## Discussion

### Repetitive elements in 4nAT genome

Repetitive elements exist extensively in the genome and have important effect on genomic instability. Therefore, studying repetitive elements can help to understand the evolution of polyploid genomes. The content of repetitive elements is always a subject of major concern for many researchers. In the processes of cloning and sequencing cDNAs of the common bean, no GC tandem repeats are found (Blair *et al.* 2009). Similarly, G/C and GC tandem repeats are much less than A/T and AT tandem repeats in the allotetraploids BACs. A study shows that there is a positive correlation between the length of tandem repeats and the frequency of variation (Schug *et al.* 1998). The full BAC sequences also exhibit a plenty of large tandem repeats. Based on these results, it is apparent that lots of the TR loci in tetraploid hybrids are extremely unstable. TEs have complicated effects on the recombination of host genomes. Most of the time, they are accompanied with indels and rearrangements of large DNA (Kazazian 2004) which can lead to genomic structural changes (Christiansen *et al.* 2008). Our analysis of 4nAT BAC sequences produces similar results to these previous studies. There are widespread gene replications in polyploids and some of the TEs might be activated, resulting in genetic variations such as recombination becoming more common. TEs are unevenly distributed across the BAC sequences of 4nAT and their distributions in genomes are distinctive. For example, the density of TEs is higher in the X chromosome and the lower recombination rate regions. At the same time, TE distribution depends on the specific characteristics of chromosomes, the TEs themselves and the organisms (Rizzon *et al.* 2002; Cridland *et al.* 2013). As regulatory elements, retrotransposons have a high tendency of amplification, which can have a large influence on the genome size of plants. For instance, they occupy at least half of the genomes of wheat (Echenique *et al.* 2002) and maize (Sanmiguel and Bennetzen 1998). By contrast, transposons have relatively less influence on the genome size of plants (Kunze *et al.* 1997). Inter-genomic displacements are common in allopolyploids, which indicate that recombination occur commonly between homoeologous chromosomes, making the different sub-genomes in allopolyploids interdependent in the evolution process (Wendel 2000). Research has shown that some tandem repeats in eukaryotes are derivatives of TEs (Sharma *et al.* 2013) and furthermore, the evolutions of different genes in polyploid plants are linked to each other (Sang *et al.* 1995; Wendel *et al.* 1995). Genomic variation is thus complicated by the interplay among different gene families and repetitive elements.

### Mechanism of recombination

Homologous and homeologous recombinations of allotetraploid fish are observed in this study. Homoeologous recombination happen in DNA segments of different progenitors (model 1 and model 2 in Figure 4) that share some degree of homology and homologous recombination occur in different sequences within the same progenitor genome (model 3 and model 4 in Figure 4). There is a research (Joseph *et al.* 2008) demonstrated that homologous recombinations are the exchange of genetic information between alleles, mediated by the conserved recombinases (such as *Rad51* and *Dmc1*), which are an integral part of mitosis and meiosis and ensure the stability of karyotypes. On the other hand, there was a hypothesis that homeologous recombination can be markedly decreased due to sequence divergence (Li *et al.* 2006). Both of the homologous and homeologous recombinations can lead to novel gene combinations and generate new phenotypes. At the same time, they can destabilize karyotypes and may even result in aberrant meiotic behavior (Gaeta and Chris 2010). In some polyploids, however, recombination of homoeologous loci is required for stability (Udall *et al.* 2005). Triggers of recombination include single-strand nicks and double-strand breaks (Szostak *et al.* 1983); however, given the current data, we do not know whether the recombination hotspots could change by different triggers. During the meiotic divisions of plant cells, recombination mainly occurs between allelic sequences of homologous chromosomes (Naranjo and Corredor 2008), but not randomly along the chromosomes. In fact, the recombination hotspots are variable in different species (Anderson *et al.* 2001). In humans, the sub-telomeres of chromosomes are hotspots of inter-chromosomal recombination (Linardopoulou *et al.* 2005); however, in human cancer cells, the ribosomal RNA gene clusters are recombination hotspots (Stults *et al.* 2009). It has been reported that simple tandem repeats, transposable elements or satellite sequences could initiate recombination and generate new genes (Yang *et al.* 2008). Herein, we find similar results except the satellite sequences initiate recombination, and we demonstrate that the mechanisms by which TRs and TEs generate chimeras are different. Given large amounts of repetitive elements have been detected in many species of eukaryote, recombination between ectopic loci seems to be inevitable. The instability of repetitive elements and the intervention of recombinase both can benefit the recombination and produce the chimeras in the hybridization of allotetraploid fish.

### TRs and chimeras

Although the main driver of recombination between any two sequences is homology (Naranjo and Corredor 2008), we find that recombination do not happen most frequently among sequences with highest similarity. There is no question about the origination and evolution of TRs involves several complex mechanisms and factors, for example, slipped-strand mispairing (SSM) was proposed as the main mechanism of TRs evolution (Levinson and Gutman 1987). Under this mechanism, shorter repeats become longer and the incidence of noncontiguous SSM increases as the repetitive regions become longer and most of the TRs in 4nAT in this study are in line with this trend. According to the studies on yeast, *Drosophila* and humans, the incidence of SSM of di-nucleotide TRs is the highest and the rate of chimera related to di-nucleotide TRs in the present research is exactly the highest and this suggest that chimeras related to TRs are linked to SSM evolutionary mechanism. There is no objection to that the TRs are unstable in the genomes. During the process of DNA replication, DSB (double-strand break) is more likely occur at TR locus and then the chimera is formed through recombination between different sequences in the process of DSB repair. When both of the parental homologous sequences contain the same tandem repeat, they are conserved between each other and it’s hard to detect the chimera, so the incidence of recombination in the corresponding homologous sequence of 4nAT is the lowest.

### TEs and chimeras

TEs have a sustained impact on genome evolution; for example, they are important ectopic recombination sites (Yang *et al.* 2008). In some allopolyploid plants, recombinations induced by TEs play an important role in the reconstruction of ribosomal DNA (Pontes *et al.* 2004). Some studies have claimed that the activities of TEs have an irreplaceable role in the generation of new genes and genome rearrangements of flowering plants (Bennetzen 2005) and are the main drivers of diversity in vertebrate genome (Pontes *et al.* 2004). Retrotransposons or retrotransposon-like sequences are located in the flanking regions of a series of genes (White and Wessler 1994) and they are demonstrated to mediate recombinations to form chimeras (Yang *et al.* 2008). Research shows that even dissimilar sequences could be linked together through TEs (Shapiro 2005), which is analogous to the recombination in the allotetraploid genome. The genome of allotetraploid hybrid fish contains the genomes of both progenitors, which result in a high rate of heterozygosity of the chromosomes. Furthermore, recombination can occur between different LTRs and may lead to the loss of some internal segments (Liu and Wendel 2002) and some of the new LTRs in the allotetraploid hybrid BAC sequences may be generated by these recombinations. Additionally, a group of tyrosine-recombinase was reported to have the ability to encode DNA transposons from pathogenic fungi (Goodwin *et al.* 2003) and it’s acceptable to infer that the TE sites benefit the recombinases have a role to play. And with the help of recombinases, the TEs can promote genomic rearrangements through ectopic homologous recombination (Belancio *et al.* 2010), which may be one explanation for why the four models of chimeras at TE loci in 4nAT show no significant difference among composite patterns a, b, c and d.

## Conclusion

The presence of the DNA chimeras found in the polyploid animal derived from the hybridization is a novel trait which is probably related to the phenotypic changes of the hybrid offspring. However, little is known about the neighbor repetitive elements of the chimeras which may be linked to the occurrences of the chimeras. Herein, we first report the DNA recombinants accompanying tandem repeats (TRs) and transposable elements (TEs) in allotetraploid hybrid fish (4nAT), which demonstrate that different models of chimeras are linked to TRs and TEs. In addition to DNA recombinants, insertions and deletions of DNA segments are linked with TRs and TEs too. These results show the occurrences of the chimeras linked to TRs and TEs can reduce the genomic shock effects and favore the adaptive capacity of the allotetraploid hybrids.

## Acknowledgments

This work was supported by National Natural Science Foundation of China Grants (30930071, 91331105, 31360514, 31430088, and 31210103918), the Cooperative Innovation Center of Engineering and New Products for Developmental Biology of Hunan Province (20134486), the Construction Project of Key Discipline of Hunan Province and China, the National High Technology Research and Development Program of China (Grant No. 2011AA100403).

